# Regulatory strategies to schedule threshold crossing of protein levels at a prescribed time

**DOI:** 10.1101/2022.02.27.482184

**Authors:** César Nieto, Khem Raj Ghusinga, Abhyudai Singh

## Abstract

The timing of diverse cellular processes is based on the instant when the concentration of regulatory proteins crosses a critical threshold level. Hence, noise mechanisms inherent to these protein synthesis pathways drive statistical fluctuations in such events’ timing. How to express proteins ensuring both the threshold crossing at a prescribed time and minimal timing fluctuations? To find this optimal strategy, we formulate a model where protein molecules are synthesized in random bursts of gene activity. The burst frequency depends on the protein level creating a feedback loop, and cellular growth dilutes protein concentration between consecutive bursts. Counterintuitively, our analysis shows that positive feedback in protein production is best for minimizing variability in threshold-crossing times. We analytically predict the optimal feedback strength in terms of the dilution rate. As a corollary to our result, a no-feedback strategy emerges as the optimal strategy in the absence of dilution. We further consider other noise sources, such as randomness in either the initial condition or the threshold level, and find that in many cases, we need either strongly negative or positive feedback for precise scheduling for events.

## I. Introduction

Proper timing of molecular and cellular events is critical for a wide range of cellular processes in development, decision-making, signal transduction, coordination of responses, etc. [1]–[8]. In many cases, transcriptional response drives the timing of important events whereby the event occurs upon accumulation of a regulatory protein up to a critical threshold level [2], [9]–[22]. The inherent stochastic nature of gene-expression causes cell-to-cell variability in the time evolution of the protein levels — and consequently in the timing of cellular events — even for cells with identical genetic content and the same environmental conditions [23]– [28]. A fundamental question of interest is to understand how cells control temporal dynamics of the underlying regulatory protein to ensure precision in event timing.

Recent works have studied precise scheduling of events for gene-expression models of varying complexity, where the timing of an event is formulated as the first-passage time (FPT) for the protein level [29]–[41]. Assuming different empirical forms for the regulation of gene-expression (e.g., feedback auto-regulation, activation/repression by an upstream component, etc.), these works search for the best model parameters that minimize the noise in timing [34]– [36], [38], [39], [42], [43]. In the limiting case where protein does not degrade (or dilute), any form of feedback gives higher noise in event timing around a fixed mean time [35]. This result is robust to extrinsic disturbances [44] and several physiologically relevant variations in model parameters [35], [45], except for the case when the initial protein amount is drawn from a distribution [46]. For a protein that degrades, positive feedback from the protein level to the transcription rate suppresses noise in event timing around a fixed mean by counterpoising the effect of degradation [35], [42], [43]. These two findings suggest a relationship between the degradation rate of the protein and the strength of the positive feedback to schedule events with precision. However, the exact nature of this relationship remains to be explored.

In this work, we consider a model of gene expression based on stochastic hybrid system formalism. Protein production is assumed to occur in bursts and is modeled as a stochastic event, whereas dilution due to cell growth is modeled using a deterministic ODE [38], [47], [48]. We provide analytical results on FPT moments and compute the optimal feedback strategy that minimizes noise in FPT around a given fixed mean under different modeling assumptions. In particular, we consider a linear form of the feedback and analyze scenarios where we draw either the initial condition or the event threshold from some positive-valued distributions, representing static extrinsic noise.

## II. Preliminaries

In this section, we formulate a model of gene-expression and describe the corresponding FPT problem. We then summarize the previous results that quantify the noise in FPT and provide optimal feedback strategies that minimize the noise.

### A. Model description

Let *x*(*t*) be the concentration of the protein of interest at time *t*. We assume that the event of interest occurs when *x*(*t*) crosses the threshold level *X* for the first time (Fig. 1). The FPT *T* when the event *x* reaches a threshold *X* is defined formally as

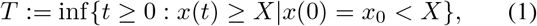

where *x*_0_ is protein level at *t* = 0.

**Fig. 1.**
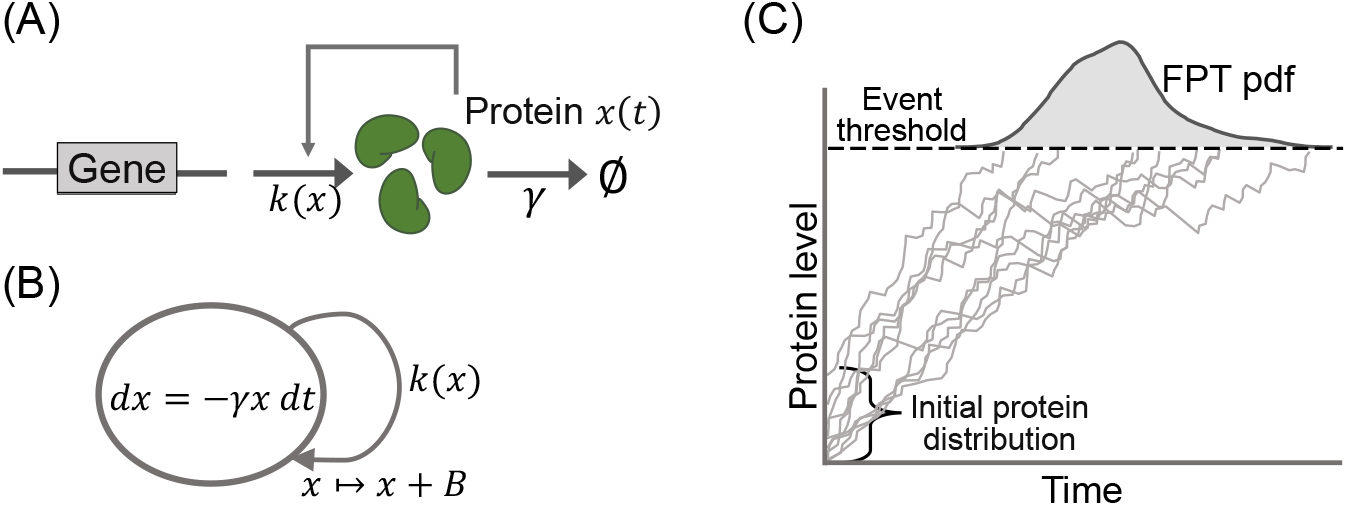
Model schematic and illustration of first-passage time. (A) Gene produces protein *x*(*t*) in bursts. The burst frequency depends upon the protein level as *k*(*x*), creating a feedback loop. The protein degrades/dilutes through a first-order kinetics with rate constant *γ*. (B) A stochastic hybrid system representation of the gene-expression model, whereby the protein level increases upon burst arrival at a rate that depends upon the protein level. Protein dilution is modeled using an ordinary differential equation. (C) First-passage time for the protein level to reach a threshold. Sample trajectories represent different realizations of the protein evolution obtained using Monte Carlo simulations. The initial protein level is drawn from a distribution.

We model the dynamics of *x*(*t*) using a stochastic hybrid system (SHS) formalism comprising both discrete jumps and continuous dilution. In particular, we assume that *x*(*t*) is produced in bursts as

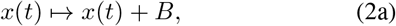

where the burst size, *B*, follows a positive-valued distribution. More precisely, the probability, ℙ, of arrival of a burst in an infinitesimal time interval (*t, t* + *dt*) is given by

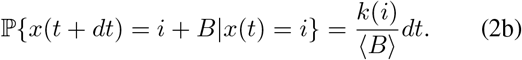

Here, we assume that the protein production rate is a function of the protein level as *k*(*x*). This is motivated by investigating how different feedback mechanisms might affect the noise in event timing. In this framework, an open-loop production (no feedback) implies a constant rate *k*(*x*) = *k*, a negative feedback means that *k*(*x*) decreases with *x*, and positive feedback signifies that *k*(*x*) increases with *x*. We have used the mean burst size ⟨*B* ⟩ as a normalization constant. Wherever needed for analytical tractability, we will assume a linear form of the feedback as *k*(*x*) = *k*_0_ + *k*_0_*x*. Here *k*_0_ is the basal rate of protein production, *k*_1_ is the strength of the feedback, and ⟨*B* ⟩ is the mean burst size used as a normalizing factor convenience.

In between successive bursts, the protein concentration is diluted by cell growth as

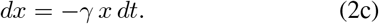

Here *γ* denotes the dilution rate. Finally, we assume that both the initial protein level *x*(0) = *x*_0_ and the threshold for the event *X* are drawn from arbitrary positive-valued distributions.

We base this model formulation on the following key assumptions. First, the protein does not degrade but only dilutes due to cell growth. This assumption itself requires that the cell volume grows exponentially over time and that the production rate in terms of protein numbers scales with the cell volume [49]–[52]. Second, both the timescale of the promoter switching and the mRNA half-life are much smaller than the turnover rate of the protein. This allows us to approximate the randomness in the dynamics of promoter and mRNA in a burst parameter. Often, the burst size follows a geometric distribution [53], but other distributions are also possible [45], [54]–[56].

The statistics of the first-passage time (FPT) defined in Eq. (1) are analytically intractable in general. However, in a limiting case where dilution is ignored, the FPT moments were computed in previous work [35]. Next, we reproduce these results and the computation of optimal feedback strategy to minimize the noise in timing from them.

### B. FPT statistics in absence of dilution

Let ℙ(*x*(*t*) = *i*) = *p*_*i*_(*t*), then in absence of dilution, i.e., *γ* = 0, the time evolution of *p*_*i*_ for Eq. (2b) is governed by the chemical master equation [57], [58]

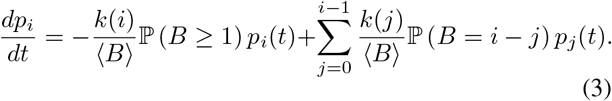

To compute the statistics of the first-passage time, *T*, as defined in Eq. (1), we note that [35]

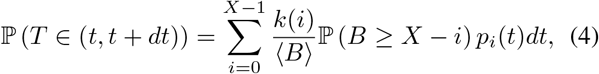

resulting in the following probability density function (pdf) of FPT

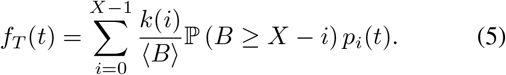

Intuitively, the process *x*(*t*) crosses the threshold *X* for the first time at time *t* + *dt* if the protein count was equal *i* at time *t* and a burst of size greater than or equal to *X* −*i* occurred in the next infinitesimal time interval (*t, t* + *dt*).

A compact form of the FPT pdf above is

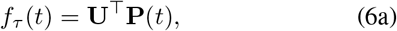

where

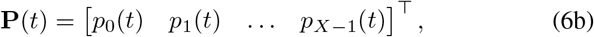

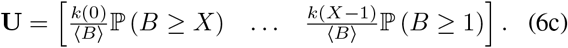

The time evolution of **P**(*t*) can be obtained from Eq. (3) as the dynamical system

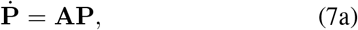

whose solution is given by

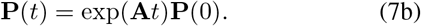

Here the triangular matrix *A* consists of the elements

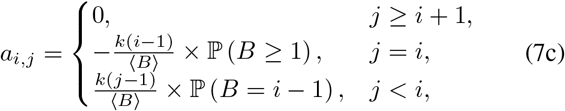

and **P**(0) is the vector of consisting of probabilities of the initial protein count, *x*_0_. Using (7b) in (6a) yields the following for the first-passage time probability density function

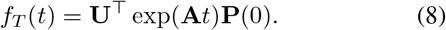

We can use Eq. (8) to compute the moments of FPT as follows

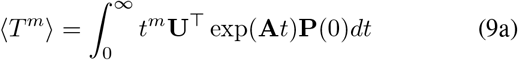

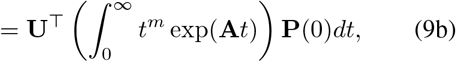

where ⟨·⟩ denotes expectation of its argument and *m* is the order of the moment. The above integral may be written as

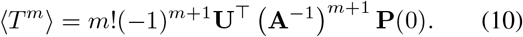

provided **A** is Hurwitz stable. This indeed the case, as each diagonal element of **A** is negative and is greater in magnitude than sum of all other elements of that column [35]. The first two moments of FPT for the special case of deterministic initial condition *x*_0_ = 0, deterministic threshold *X*, and geometrically distributed burst size are given by:

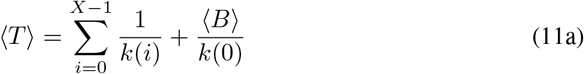

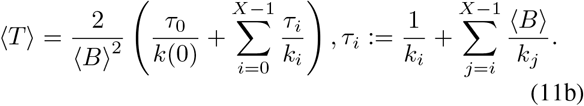

One can use conditioning argument to compute these moments when the initial condition *x*_0_ ≠ 0 and is instead drawn from a distribution [46]. Likewise, the event threshold may also be drawn from any positive-valued distribution. These scenarios are studied in section IV (Figs. 4 and 5).

### C. Optimal feedback strategy

The problem of finding an optimal feedback strategy that minimizes the noise in timing (quantified by the coefficient of variation squared = variance/mean^2^), given a fixed mean ⟨*T*⟩, is a constrained optimization problem. It yields the following analytical solution via the method of Lagrange multiplier [35]:

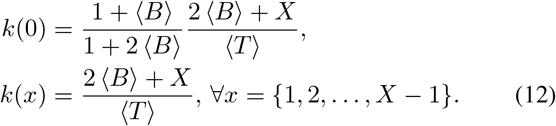

In the limit of a small burst size, the above solution is essentially a no feedback strategy. Even for a geometrically distributed burst size, all the production rates are equal except for the first one; thus, the optimal strategy is *approximately* a no feedback.

## III. Optimal feedback strategy for bursty gene expression

We next focus our attention on the case *γ* ≠ 0 and particularly restrict ourselves to the class of linear feedback, where the burst frequency takes the form (*k*_0_ + *k*_1_*x*(*t*))*/*⟨*B*⟩.

As a consequence of this linearity, the time evolution of the statistical moment of *x*(*t*) can be obtained exactly via

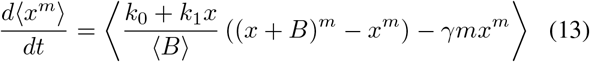

for *m* ∈ {1, 2, … } [61]–[63]. Setting *m* = 1 and assuming *x*(0) = 0 with probability one, yields the mean concentration dynamics

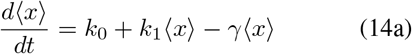

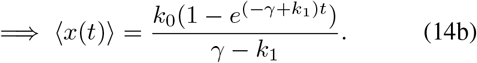

If the feedback strength *k*_1_ *< γ* then lim_*t*→∞_⟨*x*(*t*)⟩ = *k*_0_*/*(*γ* − *k*_1_), and we assume that the critical concentration threshold *X < k*_0_*/*(*γ* − *k*_1_) needed for event timing is below this steady-state mean level. In contrast, if *k*_1_ ≥ *γ* then the mean concentration grows unboundedly lim_*t*→∞_ ⟨*x*(*t*)⟩ = ∞over time. Considering small concentration fluctuations around the mean trajectory, the mean first passage time

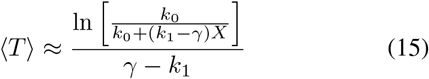

can be obtained by solving ⟨*x*(*t*)⟩ = *X*. Recall that we would like to obtain the optimal feedback strategy (i.e., the value of *k*_1_) that minimizes the noise in *T* for a given fixed mean FPT ⟨*T*⟩. Having an approximate analytical formula for ⟨*T*⟩ is quite useful in that regard as it provides the corresponding value of *k*_0_

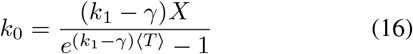

that is needed to ensure a fixed ⟨*T*⟩ as we vary *k*_1_ to explore different feedbacks.

Having set up the mean concentration dynamics we now investigate its variance by taking *m* = 2 in (13)

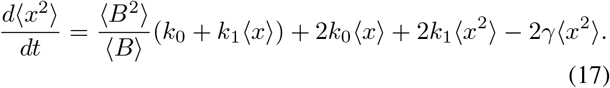

A geometric approach of connecting the variance in protein concentrations to the variance of threshold-crossing times was suggested in [64]. In particular, the variance in *T* is approximated as

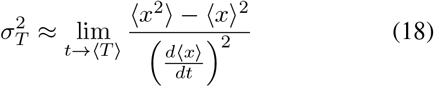

and is inversely proportional to the slope of the mean trajectory at *t* = ⟨*T*⟩ with a “flatter” approach to the threshold amplifying noise in threshold-hitting times. Substituting the solutions of (14a) and (17) in (18) results in the following analytical expression for the noise in *T* as quantified by its coefficient of variation

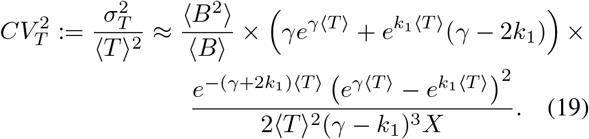

In the limit of no dilution and no feedback (*γ* → 0 and *k*_1_ → 0), the formulas (15) and (19) become exact and reduce to

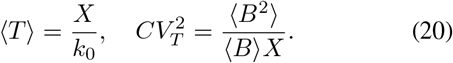

Fig. 2 illustrates the shape of 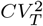 as a function of the feedback strength *k*_1_, while correspondingly changing *k*_0_ as per (16) to ensure a fixed ⟨*T*⟩. Two distinct observations can be made from this plot:

- The noise in *T* as predicted by (19) matches well with corresponding FPT noise levels obtained from stochastic simulation runs of the system (2). As expected, the match is perfect at low noise levels and begins to deviate with increasing *CV*_*T*_.
- *CV*_*T*_ first decreases with increasing *k*_1_ to reach a minimum, and then increases with increasing *k*_1_.

**Fig. 2.**
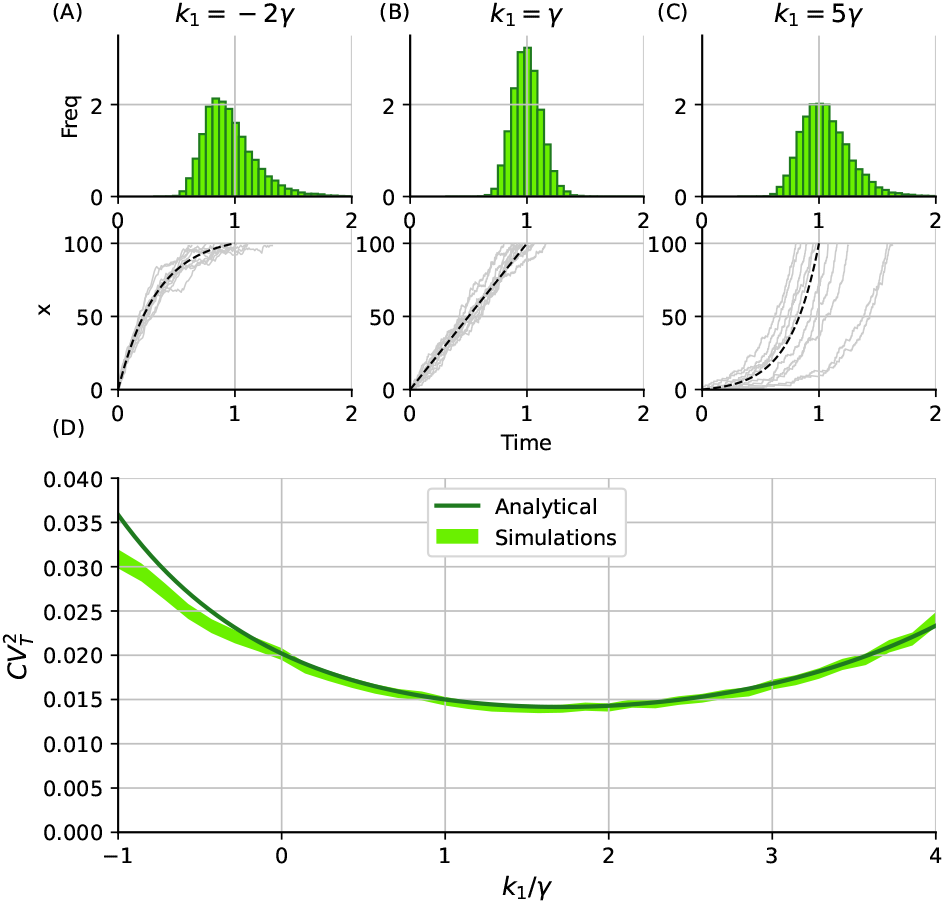
FPT fluctuations for different feedback strategies with trajectories modelled using a birth death process. The feedback strategies are classified depending on the ratio between the decay rate *γ* and the feedback strength *k*_1_. (A) *k*_1_ *< γ* (B) *k*_1_ = *γ* (C) *k*_1_ *> γ* Top: FPT histogram. Bottom: Some illustrative trajectories. (D) Noise in FPT measured by the squared coefficient on variation 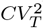 vs the feedback strength *k*_1_. Results of simulations using Gillespie algorithm (region width representing 95% confidence interval calculated with ten thousand simulation replicas) are compared to the analytical expression (19). (*γ* = 1, ⟨*T*⟩ ≈ 1, *x*(0) = 0, *X* = 100, and ⟨*B*⟩ = 1 with probability one). [59].

By solving the equation

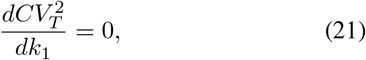

our analysis predicts the following optimal feedback strength

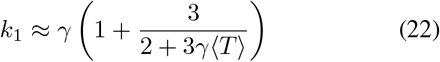

that minimizes the fluctuations in the threshold-crossing times around ⟨*T*⟩. Notice this optimal solution corresponds to a positive feedback *k*_1_ *>* 0 and depends on the dimensionless factor *γ*⟨*T*⟩. If *γ* is the exponential growth in cell volume within a cell cycle, then *γ*⟨*T*⟩ can be interpreted as mean FPT normalised by the cell-cycle duration. In the limit, *γ* → 0 a no feedback strategy emerges optimal. Moreover, the optimal feedback strength *k*_1_ ≈5*γ/*2 for *γ* ⟨*T*⟩ ≈ 0 (i.e., the mean FPT is much shorter compared to the cell-cycle duration), *k*_1_ ≈ 8*γ/*3 for *γ* ⟨*T*⟩= 1 and *k*_1_ ≈*γ* for *γ* ⟨*T*⟩ ≫1.

Does this optimal feedback strategy depend on how noise is incorporated into the model? To address these questions, we consider a simple stochastic differential equation (SDE) formulation

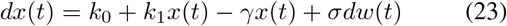

where *w*(*t*) denotes a Wiener process. Note that the mean dynamics ⟨*x*(*t*)⟩ in (23) is identical to that of the previous model and given by (14a), but now the variance evolves as

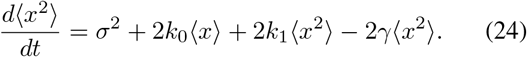

Performing the exact same analysis as before by substituting ⟨*x*(*t*)⟩ and the solution of (24) in (18) yields

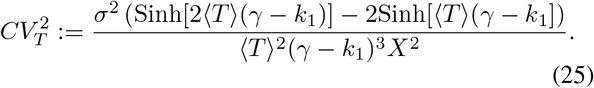

for the SDE model. While the qualitative trend of *CV*_*T*_ varying non-monotonically with respect to *k*_1_ is also seen here (Fig. 3), in contrast to the earlier results, the noise in FPT is always minimized at *k*_1_ = *γ* in the SDE model. Thus, while positive feedback provides precision in event timing in both formulations, the optimal feedback strength depends on the noise structure.

**Fig. 3.**
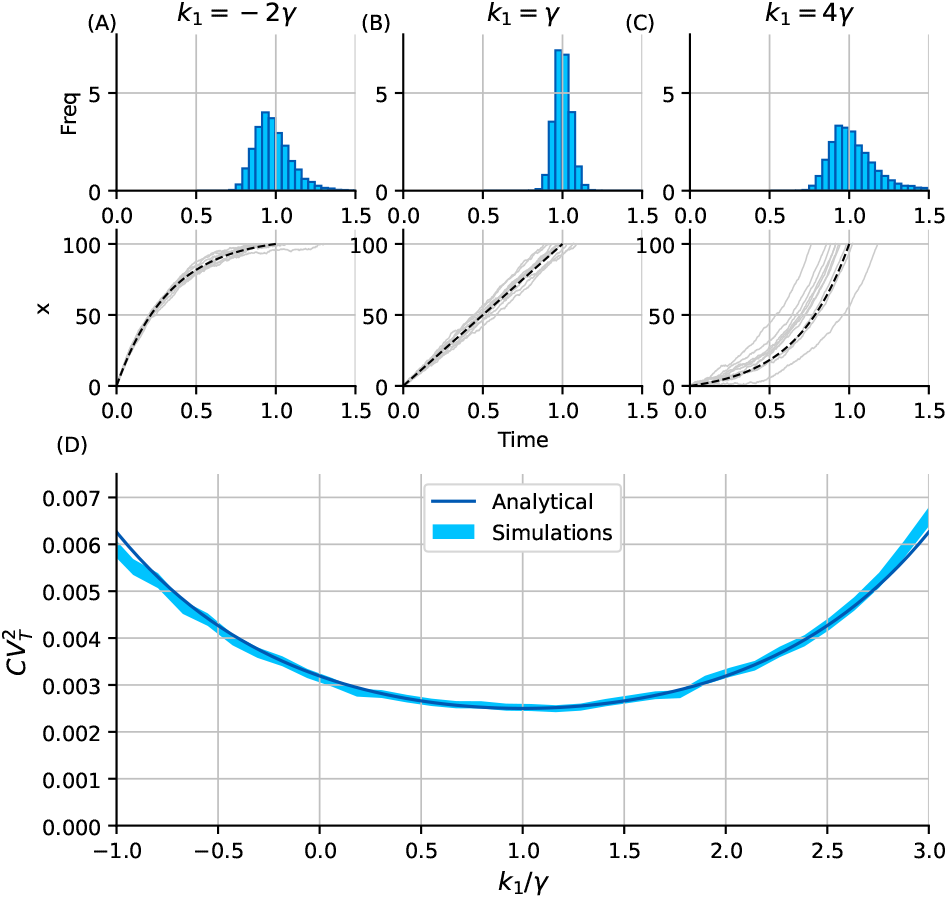
FPT fluctuations for different feedback strategies with trajectories modelled using a stochastic differential equation. The feedback strategies are classified depending on the ratio between the decay rate *γ* and the feedback strength *k*_1_. (A) *k*_1_ *< γ* (B) *k*_1_ = *γ* (C) *k*_1_ *> γ* Top: FPT histogram. Bottom: Some illustrative trajectories. (D) Noise in FPT measured by the squared coefficient on variation 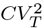 vs the feedback strength *k*_1_. Results of simulations using Euler-Maruyama algorithm [60] (region width representing 95% confidence interval over ten thousand simulation replicas) are compared to the analytical expression (25). (*γ* = 1, ⟨*T*⟩ ≈ 1, *x*(0) = 0, *X* = 100, *σ* = 5).

## IV. Optimal feedback strategy for noisy initial conditions and threshold

Our analysis in the previous section considers the synthesis of a gene product in random bursts as the predominant source of noise driving fluctuations in threshold-crossing times. We now extend this analysis to scenarios where the protein concentrations start from a non-zero initial condition *x*_0_ and build over time to the threshold *X*, where both *x*_0_ and *X* are random variables that are drawn from arbitrary distributions before each simulation run. Based on an analysis similar to (15), the mean FPT conditioned on *x*_0_ and *X* is given by

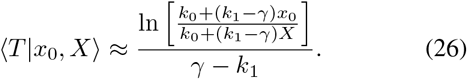

In the case of no feedback and no decay (*γ* →0 and *k*_1_ →0), this reduces to

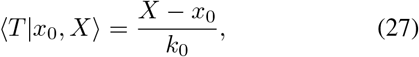

where the initial condition gets absorbed (subtracted) in the threshold. Assuming small variations in *x*_0_ and *X* around their respective means, ⟨*x*_0_⟩ and ⟨*X*⟩ results in the mean FPT

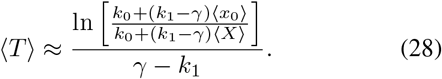

Solving this equation for *k*_0_

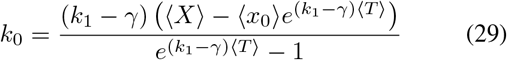

provides the corresponding changes in *k*_0_ needed to ensure a given mean FPT as we alter feedback strategies.

Our assumption of small fluctuations in *x*_0_ and *X* allows us to Taylor expand (26) around their respective means

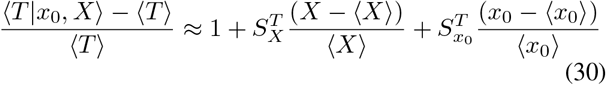

where

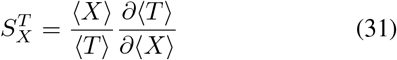

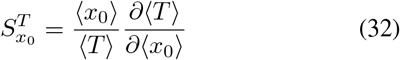

are the dimensionless log-sensitivities of ⟨*T*⟩ with respect to ⟨*x*_0_⟩ and ⟨*X*⟩, respectively. Squaring both sides and taking the expectation with respect to *x*_0_ and *X* yields the following noise in the FPTs

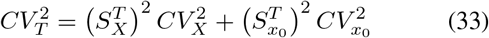

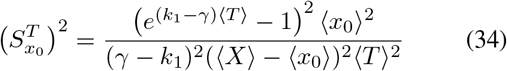

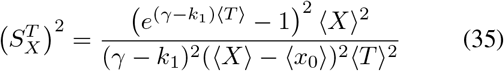

with 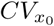 and *CV*_*X*_ being the coefficient of variation of the initial condition and threshold. Both sensitivities look similar in form but they do differ from each other in the sign of *γ* − *k*_1_ in the numerator that makes a qualitative difference – while 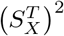 is a decreasing function of *k*_1_ with 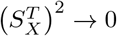 as 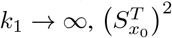 is an increasing function of *k*_1_ and 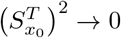 as *k*_1_ → − ∞. An important conclusion from this is that in the presence of noisy initial conditions, choosing *k*_1_ to be as negative as possible (strong negative feedback) is the optimal strategy to minimize 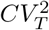 around a given ⟨*T*⟩. In contrast, for a noisy threshold choosing *k*_1_ to be as positive as possible (strong positive feedback) is the optimal strategy. We illustrate these results in Fig. 4 (a fixed threshold and a noisy initial condition) and Fig. 5 (a zero initial condition and a noisy threshold), where analytically predicted 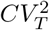 match well with those obtained from simulations confirming our predictions for different forms of feedback.

**Fig. 4.**
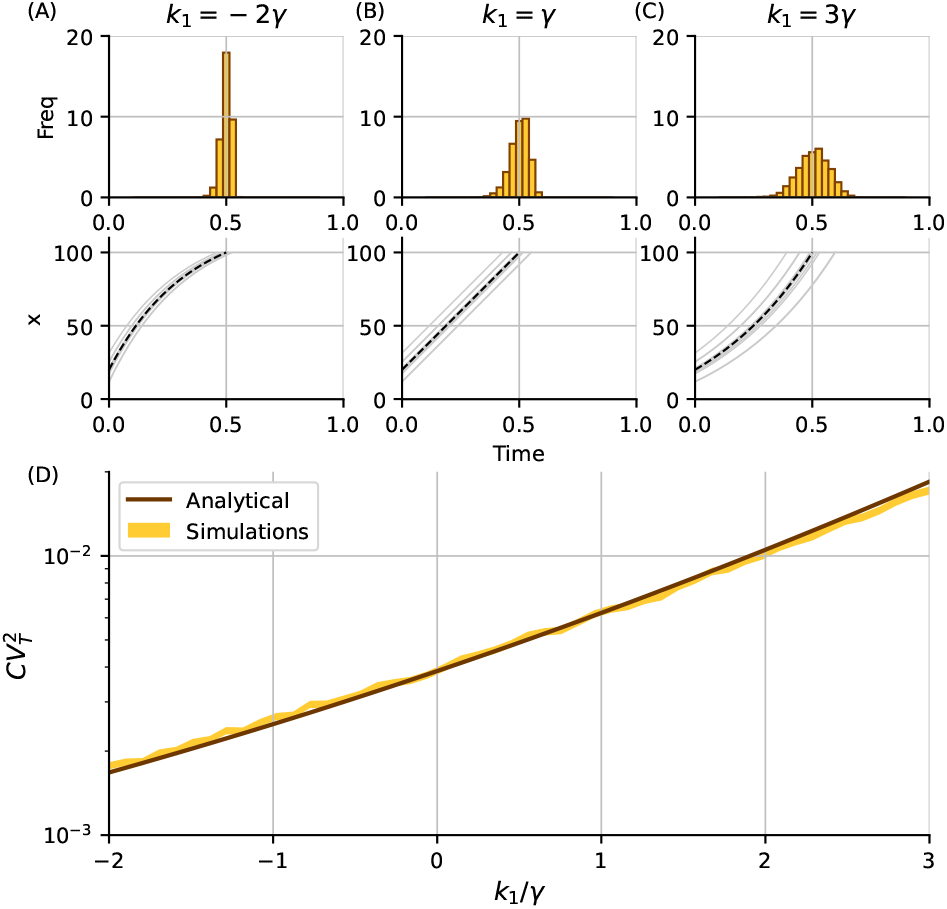
FPT fluctuations for different feedback strategies considering both random initial concentration *x*_0_ and deterministic trajectories. (A) *k*_1_ *< γ* (B) *k*_1_ = *γ* (C) *k*_1_ *> γ* Top: FPT histogram. Bottom: Some illustrative trajectories. (D) Noise in FPT measured by the squared coefficient on variation 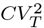 vs the feedback strength *k*_1_ relative to the decay rate *γ*. Results of simulations (region width representing 95% confidence interval for ten thousand simulation replicas) are compared to the analytical expression (34). (*γ* = 1, ⟨*T*⟩ ≈ 0.5, *X* = 100, *x*_0_ is Gamma-distributed with parameters: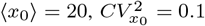).

**Fig. 5.**
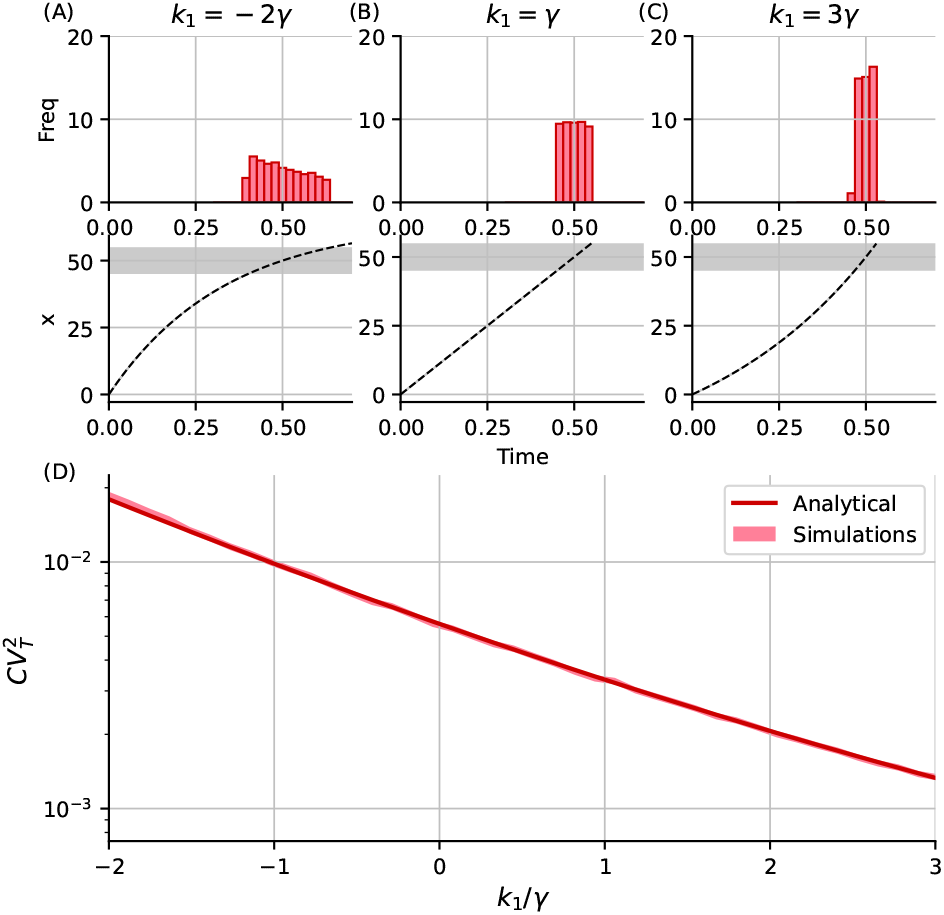
FPT fluctuations for different feedback strategies considering both random threshold concentration *X* and deterministic trajectories. (A) *k*_1_ *< γ* (B) *k*_1_ = *γ* (C) *k*_1_ *> γ* Top: FPT histogram. Bottom: Some illustrative trajectories. (D) Noise in FPT measured by the squared coefficient on variation 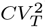 vs the feedback strength *k*_1_ relative to the decay rate *γ*. Results of simulations (region width representing 95% confidence interval for ten thousand simulation replicas) are compared to the analytical expression (35). (*γ* = 1, ⟨*T*⟩ ≈ 0.5, *x*_0_ = 0, *X* is uniformly distributed *U*(45, 55) such as ⟨*X*⟩ = 50 and 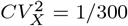).

## V. Conclusion

Uncovering mechanisms regulating the precise temporal triggering of events is vital for diverse cellular processes from development to cell-cycle regulation [1]–[8]. This contribution has explored feedback strategies that buffer fluctuations in the first-passage time around a given mean event time. To provide some biological context to this problem, consider *E. coli* cells infected by the virus bacteriophage lambda, where lysis of individual cells is the result of expression and accumulation of a single viral protein (holin) in the bacterial cell membrane up to a critical threshold [11], [12], [31], [35]. Since there is an optimal time to lyse the cells [65]–[67], the holin accumulation needs to occur to reach the threshold at the optimal lysis time. Indeed, recent experiments show that faster or slower lysis than this optimal time can result in significant fitness defects for the virus.

Using a geometric approach that connects the variance in protein concentrations to the variance in the threshold-crossing times, we derived analytical expressions for the noise in FPT’s assuming linear feedback regulation of gene expression. Our results show good agreement with Monte Carlo simulations (Figs. 2 and 3) and determined the optimal positive feedback strength needed for precision in timing. This feedback strength decreases with decreasing dilution rate and converges to a no-feedback strategy in the case of no protein decay. An SDE formulation of the problem with identical dynamics for the mean protein concentration resulted in a different positive feedback strength suggesting that this value is dependent on how noise is formulated in the model. Finally, we also considered cell-to-cell variation in FPT arising from noise in initial conditions and timing threshold in which case a strong negative or positive feedback is needed, respectively, to minimize the FPT fluctuations

As part of future work, we will consider other known sources of stochasticity, such as noise arising in the partitioning of molecules during cell division [68], [69], and extrinsic fluctuations in the growth of cell size that would be reflected in the dilution rate [70], [71]. While this work restricts the analysis to linear feedback, we will also consider Hill function type nonlinear feedback in the future, where we can explore optimal strategies by employing a combination of approximate closure schemes [62], [72], [73] and stochastic simulations.

## ACKNOWLEDGMENT

AS is supported by NIH 1R01GM124446-01.

## Notes

### Competing Interest Statement

The authors have declared no competing interest.

